# Synergizing exchangeable fluorophore labels for multi-target STED microscopy

**DOI:** 10.1101/2022.07.02.498450

**Authors:** M. Glogger, D. Wang, J. Kompa, A. Balakrishnan, J. Hiblot, H.D. Barth, K. Johnsson, M. Heilemann

## Abstract

Investigating the interplay of cellular proteins with optical microscopy requires multi-target labeling. Spectral multiplexing using high-affinity or covalent labels is limited in the number of fluorophores that can be discriminated in a single imaging experiment. Advanced microscopy methods such as STED microscopy additionally demand balanced excitation, depletion and emission wavelengths for all fluorophores, further reducing multiplexing capabilities. Non-covalent, weak-affinity labels bypass this “spectral barrier” through label exchange and sequential imaging of different targets. Here, we combine exchangeable HaloTag ligands, weak-affinity DNA hybridization and hydrophophic and protein-peptide interactions to increase labeling flexibility and demonstrate 6-target STED microscopy in single cells. We further show that exchangeable labels reduce photobleaching, facilitate long acquisition times and multi-color live-cell and high-fidelity 3D STED microscopy. The synergy of different types of exchangeable labels increase the multiplexing capabilities in fluorescence microscopy, and by that, the information content of microscopy images.

## Main

Stimulated emission depletion (STED) microscopy is a super-resolution microscopy technique with theoretically unlimited spatial resolution^1^. In combination with well-established fluorescence labeling protocols, STED is a powerful tool in optical structural cell biology. As many high-performance microscopy methods, STED microscopy demands for comparably high laser intensities, which may lead to photo-destruction of covalently bound fluorophores and subsequent signal loss. As a consequence of photobleaching, long acquisition times in live-cell experiments as well as imaging thick samples in 3D with high fidelity is challenging. Attempts to reduce fluorophore photobleaching include adaptive illumination schemes^2^, the development of photostable fluorophores^3^ or the application of fluorescent proteins with multiple off-states^4^. In addition, the number of cellular structures and proteins that can be imaged within the same cell is limited due to spectral overlap of fluorophore absorption and emission spectra^5^. Both challenges can be addressed with weak-affinity, non-covalent fluorophore labels that transiently bind to a target structure and are continuously replaced by intact fluorophores from a large buffer reservoir. These “exchangeable” labels provide an elegant way to bypass photobleaching and facilitate multiplexing and 3D imaging^6–8^. Originally introduced in the single-molecule imaging method termed point accumulation for imaging in nanoscale topography (PAINT^9^) using the hydrophobic dye Nile Red, the concept was generalized by the application of specific fluorescent ligands (uPAINT^10^) and was further extended to weak-affinity DNA-DNA hybridization (DNA-PAINT^11,12,13^), peptide-fragments (IRIS^14^), peptide-peptide interactions^15^ or protein-targeting oligonucleotides (aptamers^16^). HaloTag7 (HT7) is a self-labeling protein that covalently reacts with biorthogonal ligands (HTLs^17^). Recently introduced exchangeable HT7 ligands (xHTLs^18^) are cell membrane permeable fluorescent ligands for STED microscopy, representing a powerful alternative for live-cell STED studies.

Here we present sequential multi-method labeling combining the concepts of DNA-PAINT, PAINT/IRIS and HaloTag-based PAINT (HT-PAINT), and image six different structures in a single eukaryotic cell using STED microscopy. We report imaging cellular structures with high spatial resolution and demonstrate reduced photobleaching compared to covalent labels. We further demonstrate the applicability of our cross-method labeling approach for multi-target live-cell and 3D imaging.

We first tested various weak-affinity, exchangeable fluorophore labels by targeting and imaging different cellular structures individually with confocal laser scanning microscopy (CLSM) and STED microscopy, including xHTLs, PAINT/IRIS labels and DNA-PAINT labels. Sufficiently high ligand concentrations (100-500 nM) in the imaging buffer ensured a saturation of targets and thus quasi-permanent labeling and allowed imaging various structures with high quality (**Figure 1A**). Imaging with super-resolution STED microscopy visualized structural features obscured in diffraction-limited confocal microscopy (**SI Figure 1**). We next evaluated the applicability of our approach in confocal time-course measurements. DNA-PAINT and PAINT labels were previously shown to reduce photobleaching and facilitate time-lapse and volumetric imaging in CLSM and STED configurations^7,19^. To evaluate whether this applies to xHTLs, we determined the degree of photobleaching using time-lapse confocal microscopy of endogenously tagged vimentin-HT7 in U2OS cells^20^. We compared the HT7 covalent labeling (JF_635_-HTL) to its fluorogenic xHTL counterpart (JF_635_-HSAm^18^) at two different irradiation intensities (**Figure 1B**). For this purpose, we applied an analysis pipeline that generates a binary mask on drift-corrected time-series for image segmentation and recorded signal and background over time (^7^, Method section). Intensity-time traces showed that the signal intensity decreased strongly for the covalent labels over the first 25 consecutive frames, whereas we observed a moderate degree of photobleaching for exchangeable labels (**Fig 1B, SI Figure 2A**). These findings are in good agreement with intensity-time trace analysis for DNA-PAINT and PAINT labels (**SI Figure 2B**) and demonstrate the applicability of xHTLs for multi-frame time-lapse confocal microscopy. Next, for DNA-PAINT labels, we reduced the DNA-DNA hybridization time by supplementing the imaging buffer with ethylene carbonate (as demonstrated in DNA-PAINT-ERS^21^). We measured a further reduced degree of photobleaching (**SI Figure 2B**) possibly caused by faster replacement of photodamaged labels. We imagine that faster exchange rates for xHTLs labels could have similar effects.

**Figure 1.**
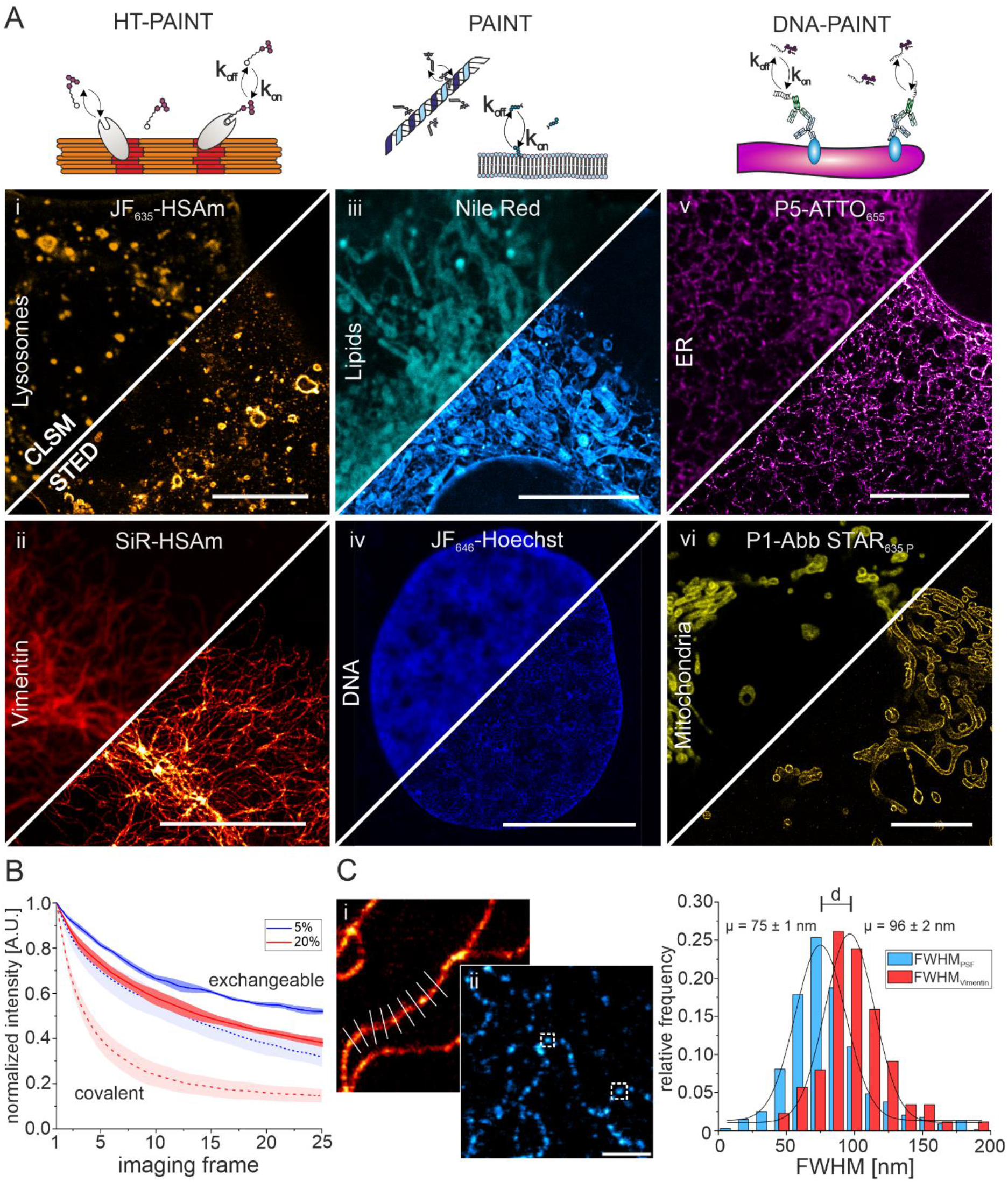
Exchangeable fluorescent labels for CLSM and STED microscopy of various cellular structures.**A**) Illustration of the principle of transient ligand binding to target structures in HT-PAINT, PAINT and DNA-PAINT (upper panel). Lower panels show CLSM and STED images using non-orthogonal xHTLs (LamP1-HT7 (i), 300 nM JF_635_-HSAm, gold; vimentin-HT7 (ii), 300 nM SiR-HSAm, red hot), PAINT labels (500 nM Nile Red (iii), cyan hot; 300 nM JF_646_-Hoechst (iv), blue)) and DNA-PAINT labels (KDEL-antibody (v), 300 nM P5-ATTO_655_, magenta hot; TOM20-antibody (vi), 300 nM P1-Abb STAR_635 P_, yellow). All scale bars in i)-vi) are 10 μm). **B**) Comparison of fluorescence signal versus time recorded in CLSM mode for vimentin structures in U2OS cells using exchangeable (JF_635_-HSAm) or covalent HaloTag ligands (JF_635_-HTL) at different laser intensities (N_cells_ = 3-5). **C**) Quantification of the resolution in STED images. The optical resolution was determined from the intensity profile perpendicular to continuous vimentin-HT7 structures (i, 500 nM SiR-C4-Sulfonamide, red hot) or the full-width half-maximum (FWHM) of single vimentin-HT7 spots (ii, 100 nM JF_635_-HSAM, cyan hot). Scale bar is 1 μm. Shown is the relative frequency distribution of the determined FWHM of single-molecule spots (light blue, n = 691) and intensity profiles (red, n = 88).

Next, we determined the spatial resolution of STED images from cells labeled with xHTLs. Making use of the inherent flexibility of transient labels to adjust the label density within the sample and control target saturation, we decided to determine the resolution in different ways. First, we used high label densities to achieve quasi-permanent labeling and imaged continuous vimentin filaments. We then determined the full-width half-maximum (FWHM) of the structure by fitting the intensity profile perpendicular to the filaments with a Gaussian function. Full labeling of vimentin filaments resulted in an average FWHM of 96 +/-2 nm (mean, s.d.) (**Figure 1C**). Second, we imaged sparsely labeled vimentin-HT7 structures followed by the determination of the FWHM of single fluorescent spots. From the distribution of single fluorescent spots, we measure an average FWHM of 75 +/-1 nm and as small as < 40 nm (**Figure 1C**). The increased average FWHM value in case of full vimentin labeling is in good agreement with our single-spot analysis considering the width of single vimentin filaments (10 nm^22^) and additional linkage error introduced by the HaloTag system (PDB-ID: 7ZJ0). We further evaluated the spatial resolution by analyzing the intensity profiles of vimentin structures at intersection points (**SI Figure 3**) and show the optical separation of single filaments closer than 86 nm. Taken together, the experiments demonstrate that xHTLs enable STED imaging with sub-diffraction spatial resolution. The main goal of this study was to demonstrate that multiple types of exchangeable labels can be combined to multi-target STED microscopy. For this purpose, we operated a standard commercial STED microscope (see Methods). Several experimental strategies were reported to improve the spatial resolution in STED microscopy^23,24,25,26^, and may be implemented in this workflow if available.

We next explored whether a combination of various types of exchangeable labels would enable multi-color imaging of multiple structures in the same cell. We first validated the cross-method labeling compatibility by sequential labeling and imaging targets with xHTLs and DNA-PAINT labels in single cells (**Figure 2A-B**). Exchange of the fluorescent labels between imaging cycles facilitated high fidelity and residual-free imaging of individual vimentin and mitochondria. Next, we explored the applicability of xHTLs for 3D STED microscopy. We first labeled chemically fixed U2OS cells expressing CalR-HT7-KDEL (endoplasmic reticulum, ER) using the xHTL SiR-HSAm and recorded z-stack images in 3D STED mode. Volumetric rendering of deconvoluted images allowed high fidelity reconstruction of the ER (**Figure 2 C, Supplementary Video 1**). Further, we used cross-method labeling and sequential STED microscopy in one spectral channel for 3D imaging of chromosomal DNA (JF_646_-Hoechst^27^) and the ER (SiR-HSAm) in the CalR-HT7-KDEL cell line. Fiducial marker-based image alignment in all dimensions and volumetric rendering allowed 2-target 3D STED microscopy (Supplementary Video 2). Encouraged by these findings, we applied our cross-method labeling approach for two-color live-cell STED microscopy. We concurrently labeled the endomembrane system and vimentin-HT7 using the membrane-permeable label Nile Red and JF_635_-HSAm, respectively. Simultaneous excitation with 561 nm and 640 nm and depletion at 775 nm facilitated two-color live-cell STED microscopy and allowed following the dynamics of both structures (**Figure 2 D, Supplementary Video 3**). Finally, we combined HT-PAINT, DNA-PAINT and PAINT/IRIS labeling methods in a single cell and achieved STED imaging of six different structures (**Figure 2E**). We used 3 far-red (SiR, JF_646_ and Abberior STAR_635 P_) and two red-emitting fluorescent dyes (AlexaFluor_594_ (AF_594_), Nile Red) and exchanged the fluorophore labels between STED imaging cycles by washing steps. Lateral shift caused by the exchange was corrected using fiducial markers. This procedure facilitated precise alignment of all imaging channels and the reconstruction of highly multiplexed STED images of a single cell.

**Figure 2.**
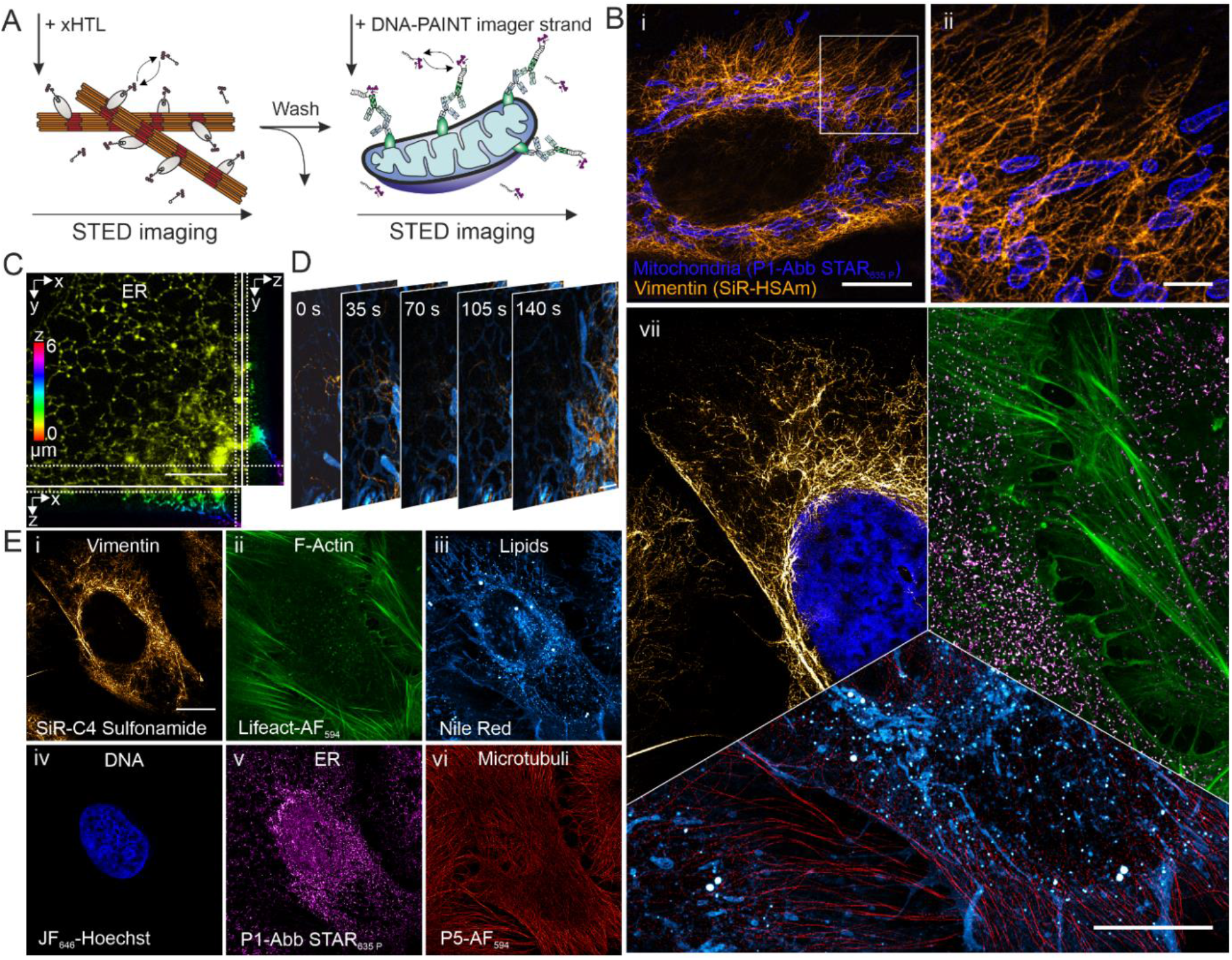
Multi-target, multi-color and multi-labeling method STED microscopy in single cells using exchangeable fluorescent labels. A) Schematic illustration of the principle of ligand exchange in multi-labeling method STED microscopy. Target specific and transient binding ligands for HT-PAINT and DNA-PAINT are exchanged between imaging rounds. B) Cross-method labeling and 2-target STED microscopy in a single cell. (i) STED image of vimentin-HT7 labeled with xHTLs (300 nM SiR-HSAm, orange hot) and mitochondria (TOM20) labeled with a DNA-PAINT imager strand (300 nM P1-Abb STAR_635 P_, blue). White box indicates magnified region shown in (ii). Scale bars are 10 μm (i) and 2 μm (ii). C) 3D-STED imaging of the ER located fusion protein CalR-HT7-KDEL in a U2OS cell. The cell was labeled using the xHTLs SiR-HSAm (300 nM). Ortho-slice shows deconvoluted, volumetric rendered 3D-STED z-stacks. Scale bar is 5 μm. D) Live-cell 2-color STED microscopy using exchangeable fluorescent and cell membrane permeable ligands. U2OS cells were labeled for the endomembrane system (500 nM Nile Red, cyan hot) and vimentin-HT7 (500 nM JF_635_-HSAm, orange hot). Scale bar is 2 μm. E) 6-target super-resolution STED image of various structures in a single U2OS cell. Cross-method labeling STED imaging was performed via HT-PAINT (i, vimentin, 300 nM SiR-C4-Sulfonamide), IRIS/PAINT (ii-iv, F-actin, 1 μM Lifeact-AF_594_, 300 nM; membranes, 300 nM Nile Red; chromosomal DNA, 300 nM JF_646_-Hoechst) and DNA-PAINT (v-vi, ER, 500 nM P1-Abb STAR_635p_; microtubule, 500 nM P5-AF_594_). Images were aligned using fiducial markers. Scale bar is 10 μm. vi) 6-target STED image in a single cell. Overlay of sequentially recorded STED images from multi-method labeling approaches (i-vi). Scale bar is 10 μm.

In summary, we demonstrate 6-target STED microscopy in single cells using exchangeable fluorophore labels. This is achieved by combining different types of exchangeable labels, employing weak-affinity DNA hybridization, exchangeable HaloTag ligands, a protein-targeting peptide, a membrane label and a DNA binder, all sharing the feature of transient target binding. We demonstrate the ability of these labeling approaches to substantially reduce photobleaching and the compatibility to STED imaging of cellular structures with sub-diffraction spatial resolution. Further improvements in spatial resolution can be achieved by tuning STED imaging parameters and ligand binding kinetics as demonstrated for DNA-hybridization^6^.

Sequential labeling with exchangeable labels is the key to enable many-target STED imaging in fixed cells, as it bypasses the limitations of covalent labels in STED microscopy caused by spectral overlap and balancing excitation, emission and depletion wavelengths. The approach is scalable, and labeling and imaging of even more targets is possible by implementing e.g. orthogonal xHTLs^18^, increasing the number of DNA-barcoded labels, by integrating orthogonal protein labels^8^, or by implementing other weak-affinity labels (^28, 29, 30^). Given the availability of highly specific target-ligand pairs in molecular biology, we envisage many more types of exchangeable labels that could turn compatible with this approach and further expand the multiplexing. Other attempts to overcome the spectral barrier in fluorescence microscopy applied fluorescence lifetime imaging (FLIM^31^), spectral unmixing^32^ or a combination of both (sFLIM^33^). These approaches can be combined with the use of exchangeable labels, and potentially help to minimize the number of washing steps by allowing pairwise imaging of labels.

Exchangeable labels continuously bind and unbind to their target, which leads to a replenishment of eventually photobleached fluorophores with intact fluorophores from the imaging buffer. This leads to an increased signal over time compared to covalent labels, which is beneficial for the image quality in 3D STED microscopy. In the case of live-cell compatible exchangeable labels, long-time live cell imaging is enabled^6^.

We envisage application of exchangeable labels for complex multi-protein studies in single cells, targeting e.g., spatial patterns of proteins and interactions in organelles at the nanoscale with high spatial resolution. Substituting covalent labels by exchangeable labels and using combinations of different labeling methods overcomes the spectral barrier and facilitates studying a multitude of proteins in the same cell. This concept will elevate fluorescence microscopy into a optical omics method for structural cell biology.

## Supporting information

Supplementary methods, figures and tables for Synergizing exchangeable fluorophore labels for multi-target STED microscopy

## Methods

Experimental procedures are described in supplementary information

## Acknowledgement

M.G., D.W., A.B., H.D.B. and M.H. gratefully acknowledge funding by the Deutsche Forschungsgemeinschaft (DFG, German Research Foundation) – SFB1177; HE 6166/17-1; INST 161/926-1 FUGG; INST 161/1020-1 FUGG. J.K., J.H. and K.J. were supported by the Max Planck Society, the Ecole Polytechnique Federale de Lausanne (EPFL) and the Deutsche Forschungsgemeinschaft (DFG, German Research Foundation), TRR 186. We thank Prof. Ivan Dikic (IBC II) for providing the cell line U2OS-Sec61b-GFP and S. Jakobs (MPINat) for providing the U2OS vimentin-HaloTag7 cells.

## Author contributions

M.H. and M.G. conceived the original idea. M.G., D.W. and A.B. carried out the experiments. J.K., J.H. and K.J. contributed cell lines, reagents, and experimental protocols for exchangeable Halo-Tag labeling and imaging experiments. M.G., D.W. and A.B. processed the experimental data and performed image analysis. M.H., M.G. and H.D.B. supervised the experimental and technical aspects of the project. M.G. wrote the manuscript with support from M.H., D.W., A.B. All authors discussed the results and finalized the manuscript.

