## Supplementary methods, figures and tables for Synergizing exchangeable fluorophore labels for multi-target STED microscopy for "Synergizing exchangeable fluorophore labels for multi-target STED microscopy"

### Experimental Methods

#### Cell Culture

Eukaryotic cells (U2OS or U2OS Vimentin-HT7) were cultured in T-75 flasks (Greiner) at 37℃ and 5% $CO_2$ in Dulbecco`s Modified Eagle Medium (DMEM) / F-12 (Gibco, Thermo Fisher, USA) containing 10% (v/v) fetal bovine serum (FBS) (Corning, USA), 1% penicillin-streptomycin (w/v) (Gibco, ThermoFisher, USA) and 1% GlutaMAX (v/v) (Gibco, USA). Cells were passaged every 3-4 days using PBS and trypsin treatment and tested for mycoplasma contamination on regular intervals. Cells used for fixed-cell imaging were seeded on fibronectin-coated (Sigma Aldrich, Germany, 0.1% (v/v) fibronectin human plasma for 30 min) microscopy chambers (µ-slides VI 0.4, Ibidi, 3 x $10^4$ cells/ channel) 24 h prior to fixation. For live-cell microscopy, cells were seeded on fibronectin-coated 8-well chamber slides (Sarstedt, Germany, 1.5 x $10^4$ cells/ well) for 24 h hours. Prior to imaging, the cells were washed with live cell imaging solution (LCIS, ThermoFisher) and temperature adjusted to avoid lateral and axial drift.

#### Plasmids

All cell lines transiently expressing HT7 fusion proteins and stable cell lines used in this study are described in detail by Kompa *et al.*[1]. For STED microscopy of lysosomes (LamP1-HT7) and the endoplasmic reticulum (CalR-HT7-KDEL) using xHTLs, cells were transiently transfected with plasmids pCDNA5/FRT/TO_TOM20-dHaloTag7_T2A_LamP1-HaloTag7 (Addgene #187078) and pCDNA5/FRT/TO_TOM20-dHaloTag7_T2A_CalR-HaloTag7-KDEL (Addgene # 187079), respectively. For this purpose, 3 x $10^4$ U2OS cells were seeded on fibronectin-coated microscopy chambers. After 24 h incubation (37℃, 5% $CO_2$), cells were transfected using Lipofectamine 3000 transfection reagent (Gibco, Thermo Fisher, USA). Briefly, 0.31 µL Lipofectamine 3000 was diluted in 10.42 µL OptiMEM medium (Gibco, Thermo Fisher, USA), and 105 ng vector DNA was diluted in 10.42 µL OptiMEM medium with 0.42 µL P3000 reagent (Gibco, Thermo Fisher, USA). Diluted DNA solution was added to Lipofectamine diluent in a 1:1 ratio and incubated for 15 min at RT. After adding the DNA-lipid complex, cells were further incubated for 48 h at 37°C and 5% $CO_2$.

#### Sample Preparation

Cell fixation and DNA-PAINT labeling

U2OS cells were chemically fixed with prewarmed (37°C) 4% PFA (Gibco, Thermo Fisher, USA) with or without 0.1% GA (Sigma-Aldrich Chemie GmbH, Germany) in PBS (Gibco, Thermo Fisher, USA) and incubated (37℃, 5% $CO_2$) for 20 min. After washing samples with PBS, cells were permeabilized and blocked for 1 h at RT using permeabilization/blocking buffer (PB, 3% IgG-free BSA, 0.1-0.2% saponin, PBS). Subsequently, primary antibodies (Supplementary Table 1) were diluted in PB, added to the chambers, and incubated for 90 min at RT. Excess primary antibody was removed by washing the sample thrice with PBS. Custom DNA docking strand-labeled secondary antibodies (Supplementary Table 1) were diluted in PB and incubated for 90 min at RT. After removing excess secondary antibodies by washing with PBS, the samples were post-fixed with 4 % PFA for 10 min at RT and finally washed thrice with PBS. Prior to CLSM and STED imaging, DNA-PAINT imager strands (Supplementary Table 2) were diluted in imaging buffer (500 mM NaCl in PBS, pH 8.3) and added to the chambers.

Cell fixation and labeling in HT-PAINT and PAINT

U2OS mother cell lines or U2OS cells expressing HT7 fusion proteins were chemically fixed with prewarmed (37°C) 4% PFA and 0.1% GA in PBS and incubated (37℃, 5% $CO_2$) for 20 min, followed by washing with PBS. Prior to CLSM and STED microscopy, exchangeable labels were diluted in PBS and added to the chambers for labeling of Vimentin, Lysosomes, ER (all using xHTLs), Lipids (Nile Red), F-Actin (LifeAct-$AF_{594}$) and chromosomal DNA ($JF_{646}$-Hoechst).

#### STED Microscopy

STED and confocal laser scanning microscopy was performed on the Abberior STED Expert Line microscope (Abberior Instruments, Göttingen, Germany) composed of an Olympus IX83 inverted microscope (Olympus, Japan) with a UPLXAPO 60x NA 1.42 oil immersion objective (Olympus, Japan), and operated by the Imspector software (v16.3.15507; Abberior Instruments, Göttingen, Germany). For fixed-cell imaging, fluorophores were excited by 561 nm laser light with a typical spectral window of 580-690 nm, or by 640 nm laser light with a typical spectral window of 650-760 nm. For simultaneous dual color live-cell imaging, two spectral windows of typically 580-630 nm and 650-760 nm were applied under 561nm and 640nm laser excitation, respectively. The stimulated emission was performed with a 775 nm pulsed laser (Abberior Instruments, Göttingen, Germany), which is far from the spectral window of the orange channel (580-690 nm, excited by 561 nm laser) compared with the red channel (650-760 nm, excited by 640 nm laser). To compensate for this, the stimulated laser intensity was increased accordingly for the orange channel to ensure imaging quality. The fluorescent emission was recorded in line sequential mode and collected on avalanche photo diodes (APDs) using a gating of 0.75ns – 8.75 ns. The pinhole was set to 0.61- 0.71 AU for STED and CLSM. The pixel size was set to 20 nm for 2D-STED microscopy and 30 nm/ 40 nm for 3D-STED microscopy. For volumetric imaging in 3D-STED mode, the z-stack step size was set to 40 nm for single color imaging and 100 nm for dual color imaging. Line accumulation was set to 10-15 and dwell times to 3-15 µs, if not stated otherwise. Exchangeable labels and applied concentrations for experiments are provided in Supplementary Table 3. Detailed imaging settings for each measurement are listed in Supplementary Table 4.

Intensity time traces of covalent and exchangeable HaloTag ligands

For intensity time trace analysis, covalent ($JF_{635}$-HTL) and exchangeable HaloTag ligands ($JF_{635}$-HSAm) were diluted to 100 nM in PBS. For covalent labeling, diluted $JF_{635}$ HTL was added to the fixed U2OS Vimentin-HT7 cells and incubated for 30 min at RT. Afterwards, cells were washed thrice with PBS. Intensity time traces were recorded in confocal laser scanning mode. 3-5 positions (approximately 10 x 10 $µm^2$) were selected for each excitation intensity and 25 consecutive frames were recorded using following imaging parameter: pixel size 20 nm, dwell time 2 µs, line accumulation 10, pinhole 0.71 AU for the xHTL $JF_{635}$-HSAm and 1.0 AU for covalent $JF_{635}$ HTL.

Multi-target fixed-cell STED microscopy

For multi-target STED imaging, fluorescent labels were exchanged between imaging rounds in sequence either by manual pipetting or assisted by a microfluidics system (Bruker, USA) controlled by Vutara's SRX software (SRX 7.0.00rc07, Bruker, Germany). Lateral shift caused by the exchange was corrected either manually or using microspheres (TetraSpeck 0.1 µm, USA) as fiducial markers. For this purpose, microspheres were diluted to 1:500 in PBS, sonicated for 10 min and added to the chambers. After settlement for 5 min, samples were washed thrice with PBS. For the microfluidic assisted exchange of labels in two-color and 6-target STED imaging, a flow rate of 600 µL/min and total volumes of 1 mL (fluorescent label) and 2.5 - 5 mL (PBS) were used for labeling and washing, respectively.

Simultaneous 2-color live-cell STED microscopy

For simultaneous 2-color live-cell STED microscopy, Nile Red and $JF_{635}$-HSAm (xHTL) were diluted in LCIS to a final concentration of 500 nM and added to living cells seeded on microscopy chambers. After temperature adjustment, 30 consecutive frames were recorded in STED mode with an interval 17s/frame. Detailed imaging settings are listed in Supplementary Table 4.

#### Image Analysis

Intensity time traces of covalent and exchangeable HaloTag ligands

Intensity time-trace analysis of covalent and exchangeable HaloTag ligands was performed using a custom-written ImageJ macro (as described by Spahn *et al.*[2]). In brief, image sequences were drift-corrected using the ImageJ plugin "*StackReg*". User defined thresholding was then used to generate binary masks for image segmentation and extraction of signal and background intensity values. After correction for edge-effects, for each frame the signals from structures were averaged, background corrected and further analyzed in Origin software (Origin2019).

Determination of spatial resolution in STED images

To determine the spatial resolution in STED images, chemically fixed vimentin-HT7 expressing U2OS cells were labeled using xHTLs at concentrations of 100 nM and 500 nM. At high label densities, continuous signal was detected along the filaments, allowing the determination of the full-width half-maximum (FWHM) of the structure. For this purpose, a custom written analysis pipeline was established in python (https://github.com/MariusGlg/Filament_width_analyzer). In brief, 10-50 lines equally spaced and perpendicular to the vimentin filaments were drawn. Intensity profiles along the line segments (n = 691) were fitted with a Gaussian function and full-width half-maximum values (FWHM) were extracted for each segment. FWHM values were then plotted as a relative frequency distribution and fitted with a Gaussian function in Origin software. At low labeling densities, the point-spread-functions (PSF) of single vimentin-HT7 spots was analyzed as a measure of the spatial resolution. For this purpose, local intensity maxima of STED images reflecting the vimentin-HT7 positions were identified in ImageJ using intensity value thresholding ("*Find Maxima*" function) based on background and signal intensities. The ImageJ plugin "*GaussFit OnSpot*" was then used for fitting Gaussian profiles onto selected positions and extracting the FWHM (n = 88). As a third measure of the spatial resolution, the intensity profiles of vimentin structures at intersection points were analyzed. In brief, intensity line profiles perpendicular to the axes of vimentin filaments that were present in close proximity to each other were analyzed using ImageJ. The measured line profiles were then fitted using a multi-peak Gaussian function in Origin software to determine their full FWHM and mean. The FWHM and the distance between the means were taken to be the physical resolution achievable with biological samples in the setup used.

3D reconstruction of STED z-stacks

For volumetric rendering of STED z-stacks, images were first background subtracted in ImageJ using a rolling ball radius of 50 pixels. Images were then deconvoluted using the ImageJ plugin "*DeconWithGaussian*" and the zposition was color-coded using the plugin “*Z-stack Depth Colorcode*” (LUT spectrum). For single-color volumetric rendering and generation of 3D movies the plugin “*3D viewer*” was used. Dual-color 3D rendering and generation of dual-color movies was conducted in Napari open source software[3].

### Supplementary Tables

#### Supplementary Table 1

Primary and secondary antibodies used for DNA-PAINT based labeling in this study.

| Antibodies | Target | Dilution | Docking strands |
|---|---|---|---|
| Rabbit-anti-KDEL, abb176333, Abcam | Endoplasmic Reticulum | 1:200 | / |
| Rabbit-anti-TOM20, sc-11415, Santa Cruz | Mitochondria outer membrane | 1:100 | / |
| Mouse-anti-α-tubulin, T5168, Sigma-Aldrich | α-tubulin | 1:200 | / |
| AffiniPure donkey-anti-rabbit, #711-005-152, Jackson Immuno-Research | Rabbit primary antibodies | 1:100 | P1 |
| AffiniPure goat-anti-mouse, #115-005-003, Jackson Immuno-Research | Mouse primary antibodies | 1:150 | P1 |
| AffiniPure goat-anti-rabbit #111-005-003, Jackson Immuno-Research | Rabbit primary antibodies | 1:150 | P5 |

#### Supplementary Table 2

DNA docking and imager strands used in this study.

| ssDNA oligo | Sequence | concentration (nM) |
|---|---|---|
| P1-docking strand | 3´-GATC TAC ATA TT-5´-antibody | / |
| P1 imager strand | 5´-TAG ATG TAT-3´-AbberiorSTAR$_{635\ P}$ | 300/500 |
| P5 docking strand | 3´-TAT GTA ACT TT-5´ -antibody | / |
| P5 imager strand | 5´-C ATA CAT TGA-3´ -AlexaFluor$_{594}$<br>5´-C ATA CAT TGA-3´ -ATTO$_{655}$ | 300/500 |

#### Supplementary Table 3

xHTLs and PAINT labels for CLSM and STED microscopy.

| Exchangeable label | Stock solution | concentration (nM) |
|---|---|---|
| $JF_{635}$-HSAm | 100 µM (DMSO) | 100, 300, 500 |
| SiR-C4-Sulfonamide | 100 µM (DMSO) | 100, 300, 500 |
| SiR-HSAm | 100 µM (DMSO) | 300 |
| $JF_{646}$-Hoechst | 100 µM (DMSO) | 300 |
| Nile Red | 100 µg/mL | 300/500 |
| LifeAct-$AF_{594}$ | 1 mM (PBS, DTT, 10% glycerol) | 1000 |
| JF635-HTL | 100 µM (DMSO) | 100 |

#### Supplementary Table 4

Supplementary Table 4 contains a detailed overview of all CLSM and STED imaging parameter and is provided as a separate .xlsx file.

### Supplementary Figures

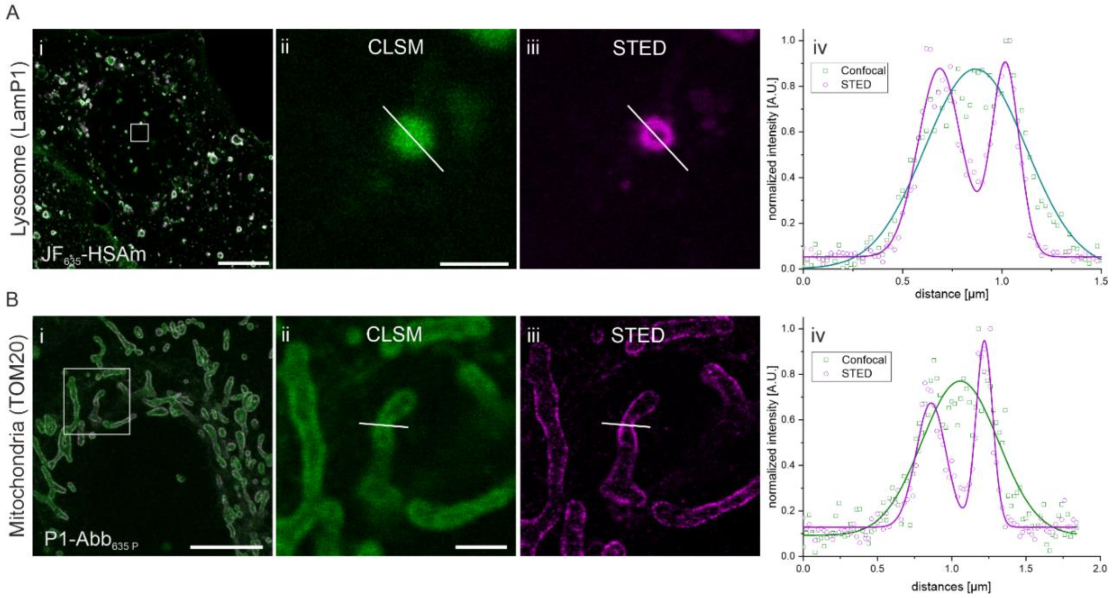

**Supplementary Fig 1.** Confocal and STED microscopy of lysosomes and mitochondria using exchangeable fluorophore labels. A) Confocal laser scanning microscopy (CLSM, green) and STED (magenta) image of lysosomes (LamP1-HT7, transient transfection) labelled using xHTLs ($JF_{635}$-HSAm) in a U2OS cell. i) Overview composite image with white box indicating a region of a single lysosome magnified in ii) and iii). iv) Normalized intensity profile across the lysosome (marked as white lines in ii) and iii)). The intensity distribution was fitted with a single (green) or two Gaussian functions (magenta). Scale bars are 10 µm (i) and 1 µm (ii-iii). B) CLSM (green) and STED (magenta) image of mitochondria (TOM20) labelled via DNA-PAINT (P1-$Abb_{635\,P}$) in a U2OS cell. i) Overview composite image with white box indicating a region magnified in ii) and iii). iv) Normalized intensity profile across single mitochondria (marked as white lines in ii) and iii). The intensity distribution was fitted with a single (green) or multi-gaussian function (magenta). Scale bars are 10 µm (i) and 2 µm (ii-iii).

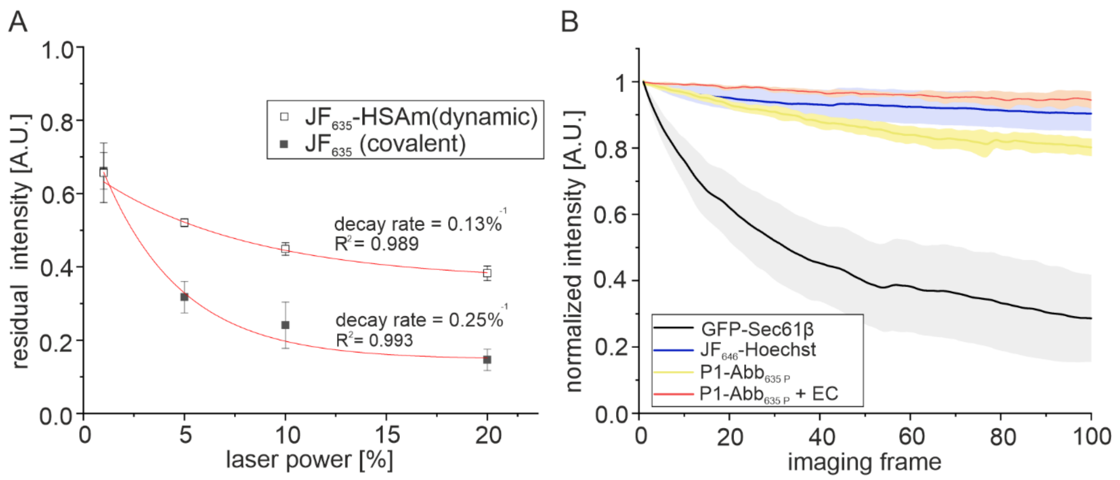

**Supplementary Figure 2**. Photostability characterization of different labelling approaches. Intensity over time trace for covalent versus transient labels (xHTLs) using confocal laser scanning microscopy. A) Quantitative analysis of the residual intensity after 25 consecutive frames and decay rates in U2OS cells expressing vimentin-HT7 and labelled with xHTLs. Comparison of covalent (JF$_{635}$-HTL) and transient (JF$_{635}$-HSAm) labels with increasing laser power (1 - 20 %). Decay rates were determined by fitting the residual intensity values with a mono-exponential decay function. Shown are mean values (N$_{cells}$ = 5) ± standard deviations. B) Intensity-time trace analysis of covalent and transient labels over 100 frames at a laser intensity of 2 %. As a covalent label the ER-located fusion construct GFP-Sec61b was constitutively expressed in U2OS (grey line) and irradiated with 488 nm. For DNA-PAINT based exchangeable labels, the ER of U2OS cells was labelled with primary antibodies against KDEL and custom DNA-docking strand labelled secondary antibodies. Confocal imaging was performed using the imager strand P1-Abb$_{635 P}$ (yellow line) at a concentration of 100 nM in PBS supplemented with 500 mM NaCl (pH 8,3). Kinetic tuning of the imager strand P1-Abb$_{635 P}$ (red line) strand was achieved by the addition of ethylene carbonate (5% (w/v)) to the imaging buffer. As an exchangeable PAINT label the chromosomal DNA stain JF$_{646}$-Hoechst (blue line) was used at a concentration of 300 nM in PBS. All images were acquired at the Leica TCS SP8 confocal microscope. Shown are mean values (N$_{cells}$ = 10) ± standard deviations. For the analysis of the intensity-time traces, a binary mask was applied to drift-corrected time-series and signal was segmented from the background. After background subtraction, intensity values were normalized with respect to the highest value.

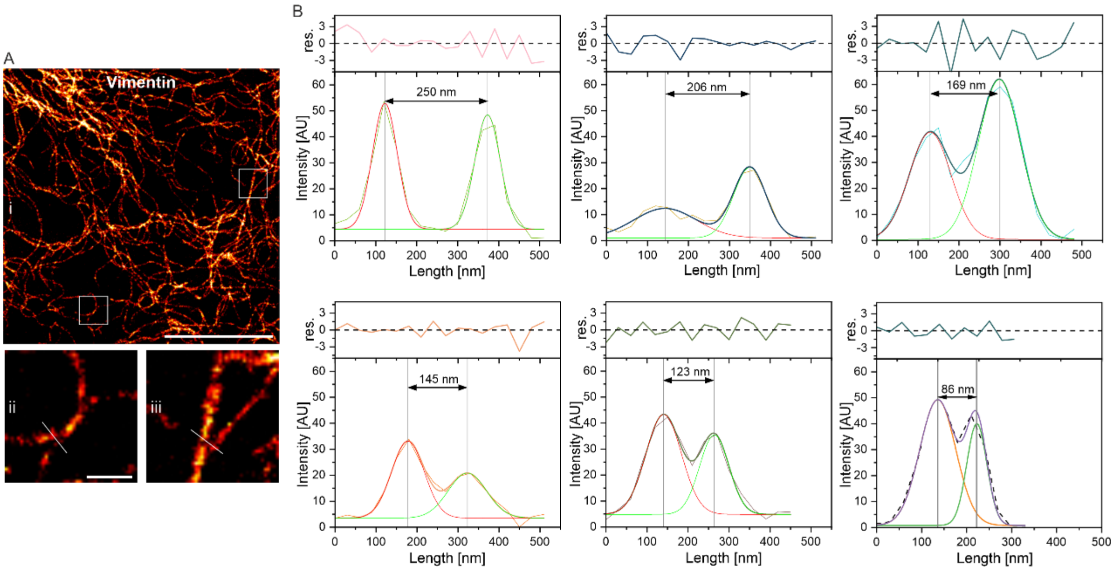

**Supplementary Figure 3**. Determination of the spatial resolution in STED images using vimentin intersection points. A) Super-resolved STED microscopy image of a U2OS cell expressing HaloTag7 conjugated to vimentin (vimentin-HT7). Vimentin-HT7 was labelled with the xHTL SiR-HSAm (500 nM in PBS). ii – iii) Expanded views of the inlays shown as white boxes in (i). White lines indicate vimentin intersection points. Scale bars are 5 µm (overview) and 500 nm (magnified regions). B) Representative intensity distributions of various positions of two vimentin intersection points. Individual intensity distributions were fitted with a gaussian function and presented with residuals of the fit. The average FWHM of vimentin intensity distributions analyzed is 76 ± 20 nm (mean ± s.d.). The narrowest center-to-center distances of vimentin at intersection points (86 nm) reflects the minimal achievable resolution in the STED images.

### Supplementary Videos

Supplementary_Video_1.mp4

Title: Single-color 3D-STED microscopy of the endoplasmic reticulum in U2OS cells labelled using xHTLs.

Legend: 3D-STED image acquisition and volumetric rendering of U2OS cells labelled for the ER-located fusion proteins CalR-HT7-KDEL using the exchangeable HT7 ligand SiR-HSAm (300 nM). The axial position is color-coded (range 6 µm) using “spectrum” LUT.

Supplementary_Video_2.avi

Title: 2-color 3D-STED microscopy in a single cell using exchangeable fluorescent ligands.

Legend: U2OS labelled for the ER (CalR-HT7-KDEL) and chromosomal DNA using the exchangeable ligands SiR-HSAm (300 nM, magenta hot) and $JF_{646}$-Hoechst (300 nM, blue), respectively. Scale bar is 5 µm.

Supplementary_Video_3.mp4

Title: 2-color live-cell STED microscopy using exchangeable fluorescent ligands

Legend: Live U2OS cells labelled for vimentin (vimentin-HT7, orange hot) and cellular membranes (light blue) using the exchangeable fluorescent labels $JF_{635}$-HSAm (500 nM) and Nile Red (500 nM), respectively. 30 consecutive frames were acquired with a framerate of 17 s.
